# 3D photogrammetry as a low cost, portable and noninvasive method for acoustic modeling of hearing

**DOI:** 10.1101/2024.09.25.614918

**Authors:** Karsten Krautwald Vesterholm, Felix T. Häfele, Florence Figeac, Lasse Jakobsen

## Abstract

1. Animals with specialized hearing such as bats utilize the directionality of their hearing for complicated tasks such as navigation and foraging. The directionality of hearing can be described through the head related transfer function (HRTF). Current state of the art for obtaining the HRTF involves either direct measurement with a microphone at the eardrum, or a μCT (micro computed tomography) scan to create a 3D model of the head for acoustic modelling. Both methods usually involve dead animals.
2. We developed a 3D photogrammetry approach to create scaled 3D models of bats with sufficient detail to simulate the HRTF using the boundary element method (BEM). We designed a setup of 28 cameras to obtain 3D models and HRTF from live awake bats. We directly compare the mesh models generated by our photogrammetry method and from μCT scans as well as the simulated HRTFs from both with measurements using an in-ear microphone.
3. Geometries of the mesh models match well between photogrammetry and μCT, but with increasing errors where line of sight is compromised for photogrammetry. The resulting HRTFs are in great agreement when comparing μCT and in-ear measurements to photogrammetry (correlation coefficients above 0.6). The 3D model and simulated HRTF of the live and awake bat likewise aligns well to the results from the deceased animals.
4. Photogrammetry is a viable alternative to μCT scans for the generation of surface models of small animals. These models allow numerical modelling of HRTFs at biologically relevant frequencies. Moreover, photogrammetry allows for model generation and subsequent HRTF simulation of live, awake animals, abolishing the need for euthanasia and anesthesia. It paves the way for large scale acquisition of 3D models for various purposes including HRTFs.

## 1. Introduction

Sound serves critical roles in communication, food finding and orientation across the animal kingdom (Bradbury & Vehrencamp, 2011). For many animals, sound source localization is crucial, and most animals localize sounds using monaural and binaural cues. Monaural cues are generated by the directionality of the pinna (if present) and diffraction of sound by the head and torso (Heffner & Heffner, 1992). Binaural cues result by comparison of the input received by two ears using the difference in arrival time of a sound at the two ears, interaural time difference (ITD), and difference in intensity received by the two ears, interaural level difference (ILD). ITDs depend on the distance between the two ears and the angle of incidence from the midline of the listeners’ head and clearly favor large animals as the ITD’s become exceedingly small with decreasing head size (Heffner & Heffner, 1992). The binaural cues contributing to the IID’s are caused by the head and pinna forming an acoustic shadow. When sound impinges the head at an angle different from zero, the acoustic shadow attenuates sounds differently on the two ears, causing a difference in sound intensity between them. The monoaural cues are pinna-specific imposed spectral characteristics to sounds reaching the eardrum. Sounds enter the pinna with different admittance, depending on their frequency, and direction in relation to the direction of the pinna which can be described by the Head-Related Transfer Function HRTF (Aytekin, et al., 2004; Firzlaff & Schuller, 2003; Firzlaff & Schuller, 2004; Fuzessery, 1996; Heffner & Heffner, 1992; Jen & Chen, 1988; Obrist, et al., 1993).

Echolocating bats navigate and forage by emitting high frequency sound pulses and localizing and identifying objects in their surroundings from the returning echoes (Griffin, et al., 1960), and they likely rely heavily on accurate sound localization to do so. Bats are generally very small, and it is therefore believed that they mainly utilize ILD’s for horizontal sound localization and monoaural directionality generated by the large pinna for vertical sound localization where elevation-depended spectral notches generated by the large tragus plays an important role (Aytekin, et al., 2004; Chiu & Moss, 2007; Lawrence & Simmons, 1982; Pollak, 1988; Wohlgemuth, et al., 2016; Wotton, et al., 1995). Thus, to understand the available cues for localization for echolocating bats and other acoustically dependent animals it is crucial to measure their HRTF.

Traditionally the HRTF is measured by inserting a microphone into a dead animal at the approximate location of the eardrum and recording sound emitted from speakers placed around the head of the animal (Guppy & Coles, 1988; Jen & Chen, 1988; Firzlaff & Schuller, 2003). While this type of experimental measurements represents the golden standard, it has several shortcomings: i) the spatial resolution of the measurements is generally limited by the practical placement of speakers and the physical dimensions of the individual components to achieve high angular resolution. ii) frequency range is limited by available hardware and the resulting system noise floor which may not cover the entire range for some species. iii) testing the influence of morphological and specialized structures (e.g. the tragus) is cumbersome, and it is not possible outside of morphological limitations. iv) the measurements require dead animals which can be difficult for rare and/or threatened species. To alleviate some of these shortcomings a more recent approach employs numerical modeling of the HRTF using high-resolution 3D models generated using μCT scanning as a viable and accurate alternative to traditional in-ear measurements (Müller, 2004; De Mey, et al., 2008). μCT scanning is a volumetric scanning technique that is not sensitive to self-occlusion and allows for very high spatial resolution and larger frequency range in the estimates. Additionally, it allows wide-scale manipulation of the 3D model to study how overall pinna morphology and orientation impact the HRTF as well as the role of specialized structures such as the tragus (Müller, 2004). However, μCT scanners are expensive, and the scanning procedure requires the animal to be completely motionless for an extended period. It is possible to use anesthetized animals, but most studies utilize dead animals as even small movements connected to breathing may compromise the 3D mesh generation.

Several scanning techniques aside from μCT, are used to generate 3D mesh models including laser scanning, magnetic resonance imaging (MRI), computed tomography (CT), and photogrammetry (Pollack, et al., 2022). But for animals, μCT scanning is still the preferred technique for creating 3D models with the purpose of simulating HRTFs (Müller, 2004; De Mey, et al., 2008; Juhl, et al., 2009; Vanderelst, et al., 2010; Gao, et al., 2011; Teshima, et al., 2022). A photography-based 3D model generation technique using jump-diffusion has been applied that allowed HRTF estimates and validation up to 3 kHz (Rebillat, et al., 2014). However, to date there is no alternative to the quality of μCT scans for accurate modelling acoustics of animals (bats, mice etc.) at high frequencies up to and beyond 100 kHz.

Photogrammetry is a non-contact surface scanning technique used to reconstruct a 3D model from a series of overlapping images. The technology is based on identifying common feature points in images and utilizing the internal parameters of the camera (focal length, sensor size, etc.) along with the relative position and orientation of the camera for the generation of a 3D model (Linder, 2009). For photogrammetry to work best, view angles of the cameras that have line of sight to the same point on the subject must be large (∼90°). Photogrammetry is inherently limited by line-of-sight and is therefore sensitive to occluded pinna geometry. Recent advances within photogrammetry have made this approach to generate 3D models highly accurate and easily accessible using cheap over-the-counter components. Photogrammetry has already been used to create 3D models of animals for a range of applications (Irschick, et al., 2022). The accuracy and ease of use make it a highly attractive alternative to μCT scans for acoustic modelling. Recent studies also suggest using photogrammetry to simulate human HRTFs, but all studies to date conclude that further work is needed to increase accuracy and repeatability (Mäkivirta, et al., 2020; Pollack, et al., 2022; Di Giusto, et al., 2023; Pollack, et al., 2023; Pollack, et al., 2023).

The aim of our study is to develop a refined photogrammetry approach to obtain 3D morphological models of bats allowing for accurate HRTF estimates at biologically relevant frequencies, up to and beyond 100 kHz, using numerical modelling. In doing so we also aim to establish a setup capable of generating sufficiently accurate 3D models of live and awake bats.

## 2. Materials and methods

We validate 3D mesh model generation using photogrammetry by comparing HRTF estimates using the Boundary Element Method (BEM) to model generation from μCT scans, and measurements from previously deceased animals using an in-ear microphone. Additionally, we evaluate how mesh model generation methods, photogrammetry and μCT, replicate a small subject with intricate details through a direct model comparison as well as their HRTF estimates using BEM. For this purpose, we used a 1:1 3D printed *Myotis daubentonii* head (Juhl, et al., 2009), a realistic subject that guaranteed to hold shape between treatments. Finally, we analyze the implications of the physical limitations in the camera setup designed for live and awake bats that are caused by the physical size of cameras and lenses.

### 2.1 Camera setup

The base camera requirements are: i) controllable focus, f-number, ISO sensitivity, and shutter speed. ii) macro photography with prime lens. iii) remote triggering (wire or wireless). We used a mix of Nikon D5600, and Canon EOS 250D, equipped with either Sigma 105mm f/2.8 EX DG OS HSM macro lens or Nikkor Micro 200 mm f/4 which fulfilled our requirements. All cameras were connected to a single computer via USB with the software CaptureGRID4 (Kuvacode Oy, Finland, Table S1) controlling camera settings and managing image download. To facilitate simultaneous triggering of the cameras, we used ESPER TriggerBox (ESPER ltd, United Kingdom). Each TriggerBox can trigger six cameras via cable, and multiple boxes can be connected through daisy chaining. All images were initially stored in native RAW format and subsequently converted to uncompressed Tag Image File Format (TIFF) using XnConvert (XnSoft, France, Table S1) with auto correction for contrast, levels and noise reduction enabled.

For the measurements of deceased animals and the 3D print (3D printer: Anycubic Photo Mono X, resin: PrimeCreator Value UV-DLP Resin), we placed seven Canon EOS 250D cameras along two vertical arcs separated by approximately 20° with all cameras pointing at a common focal point where the subject was placed (Fig. 1a). We illuminated the subject using six LED light panels (Nanlite Compac 40B, Nanguang, China). A turntable rotated the subject 360° in 5° increments, where images were captured with all seven cameras simultaneously before the turntable was rotated the next 5° (spatial image distribution shown in Fig. 1c). A total of 511 images are captured during a full rotation. The redundancy in the number of images captured allows investigation into the number of images and the angular separation between images necessary to generate a usable model and a validation of the camera setup for live animals. We used the aperture priority setting on the cameras, with f-stop f/13, ISO at 100, and the shutter speed set to ‘auto’.

**Figure 1:**
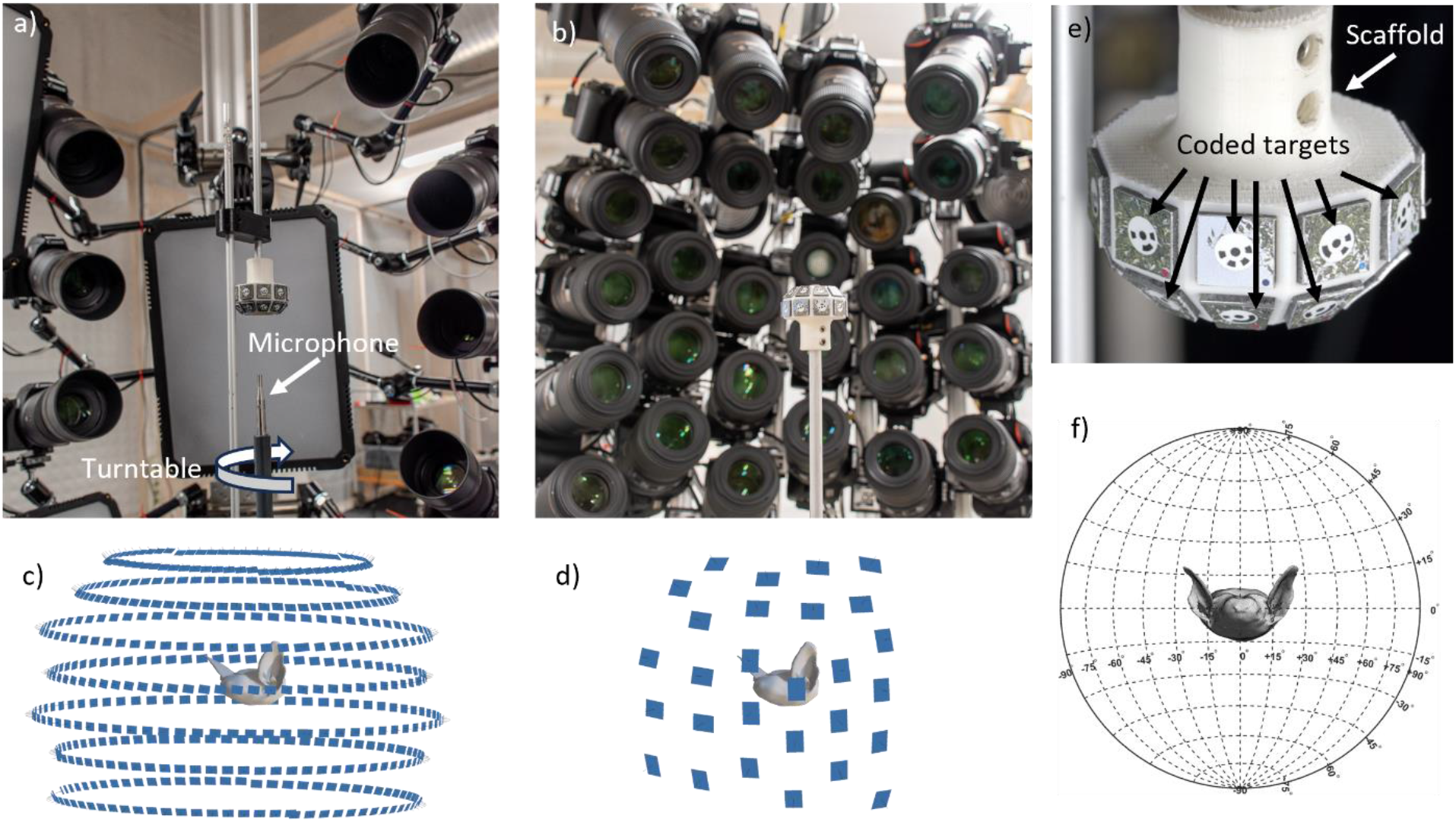
Overview of image capture setups and pre-processing steps for the resulting 3D model. a) 7-camera turntable setup for inanimate bats. b) 28-camera setup for alive bats. c) Spatial configuration of the 511 images captured by the 7-camera turntable setup. d) Spatial configuration of the 28 images captured by and the 28-camera setup for alive bats. e) Scaffold holding coded targets printed on aluminum sandwich material. f) Aligned photogrammetry model of 3D print shown in the projection used when illustrating the HRTFs.

For alive animals, we placed 28 cameras as closely spaced as possible in an aluminum frame rig, consisting of 5 variably spaced arms. The subject is covered from view angles of approximately 75° azimuth and approximately 75° elevation (Fig. 1b, spatial image distribution shown in Fig. 1d). One Nikon camera was equipped with the Nikkor Micro 200 mm f/4 lens, the remaining 27 cameras were equipped with Sigma 105mm f/2.8 EX DG OS HSM macro lenses. Lighting consisted of 3 Godox ML30 LED lights equipped with Godox FLS5 Fresnel lenses (Godox, Shenzhen, China). We used manual settings with f-stop f/14 and shutter speed 1/400 seconds and ISO set to auto on all cameras. For triggering we used a custom MATLAB (Table S1) app to control the ESPER TriggerBox for faster, synchronized image capture instead of the TriggerBoxControllerV1 accompanying the ESPER TriggerBox which we used for the seven-camera setup.

We used coded targets to aid in image alignment and to scale the 3D model to correct dimensions. A scaffold was designed to hold 20 coded targets (Fig. 1e) in a hemispherical shape for a subset of the targets to always be visible. The hemispherical shape of the scaffold increases the accuracy of the scaling of the 3D model. The feature detection-based image alignment in the photogrammetry pipeline was improved by printing a unique background with many visual features on each coded target. The coded targets were printed on aluminum sandwich material in standard printing quality. The scaffold was placed immediately below or above the subject ensuring that it was visible in all images without obscuring the subject.

### 2.2 Photogrammetry process

We used Metashape Professional (Agisoft, Russia, Table S2) to create 3D models from the captured images, utilizing the workflow and settings described by (Over, et al., 2021): i) Photo alignment and sparse cloud generation, ii) alignment optimization and scaling, iii) model generation. We adjusted the parameters as listed in Table S2 given our specific application and exported the resulting 3D model in .ply-format.

### 2.3 μCT scanning

We obtained μCT generated 3D models of two previously deceased *M. daubentonii* and the 3D printed *M. daubentonii* using a vivaCT 40 μCT scanner (SCANCO medical AG, Switzerland). The two deceased *M. daubentonii* were preserved in ethanol and dried using paper towels and a blow dryer for approximately 5-10 minutes before scanning like De May, et al. (2008). We processed the Digital Imaging and Communications in Medicine (DICOM) images from the scan into 3D models as isosurfaces using Amira (ThermoFisher, USA, Table S2).

### 2.4 Pre-processing and editing of 3D models

By following the pre-processing steps described by Brinkmann, et al. (2023), we ensured the 3D models meet the requirements for sufficiently accurate numerical calculations. To this end, we chose mesh element sizes of all 3D models suited for the highest biological relevant frequencies. For *M. daubentonii* (relevant frequencies up to 100 kHz), the mesh elements were < 0.57mm, and for *M. nattereri* (relevant frequencies up to 130 kHz) the elements were < 0.44mm (Marburg, 2002).

All 3D models in this study have the same alignment, nose pointing along the positive x-axis and the y-axis going through the eyes, and the origin place right between the eyes (Fig. 1f). This orientation of the bat is slightly different from what is required by mesh2hrtf, that states the y-axis should go through the interaural center (Brinkmann, et al., 2023), but testing both configurations showed no difference in the HRTF estimates.

For comparison of HRTFs, we manually edited the mesh in the ear canals of both the μCT scan and photogrammetry models, such that the position of the acoustic receiver was the same.

### 2.5 Acoustic modeling of the HRTF

Our acoustic modeling of the HRTF follows the pipeline for the mesh2hrtf toolbox (Ziegelwanger, et al., 2015) as presented by Brinkmann, et al (2023). Mesh2hrtf uses the boundary element method (BEM) to model the HRTF and we modelled the entire boundary as rigid following De Mey, et al. (2008) and Rebilliat, et al. (2014). Mesh2hrtf uses the principal of reciprocity, placing a source in the ear canal, and receivers in the surrounding space to speed up calculations. An acoustic source is defined in mesh2hrtf by manually assigning a velocity boundary condition to a single triangle element of the 3D model inside the ear canal. Sound pressure is then simulated in the surrounding space at designated evaluation points. The sound pressure simulated at each evaluation point is then referenced using a reference pressure associated with the specific evaluation point. When applying the principal of reciprocity, the reference pressure is defined as the pressure at the interaural center given a monopole placed at the appropriate evaluation point. This is the standard method of referencing for HRTF (Møller, 1992). The simulated sound pressure at the evaluation points only reflects the influence of the head and pinna and is therefore interpreted as the HRTF.

The frequency ranges used are 15 kHz – 130 kHz for *M. nattereri* and 25kHz – 100 kHz for *M. daubentonii* in increments of 1kHz respectively. To enable this frequency range, we changed the default frequency range in the source code of mesh2hrtf. The HRTF was evaluated on a sphere with 1 m radius, spanning elevation angles from 0° to 180° in 100 steps (1.8° increments) and azimuth angles from -180° to 178.2° in 199 steps (1.8° increments) totaling 20,000 evaluation points. The simulated HRTF was stored in Spatially Oriented Format for Acoustics (SOFA) for further processing in MATLAB. The HRTF is visualized using Lambert’s equal-area azimuth projection (Maling, 1992) (Fig. 1f). The equal-area property of this projection allows the area of the lobes to be directly compared. This comes at the cost of the introduction of minimal angular distortions, meaning the area of lobes can be compared but not the shape (Müller, 2004).

### 2.6 HRTF measurements

We measured HRTFs of previously deceased bats (one *M. daubentonii* and one *M. nattereri*) by comparing the sound field of a multi-harmonic sweep within the ear of the bat to the sound field without the bat. A total of 15 loudspeakers (Series 7000 Electrostatic Ultrasonic Sensor, SensComp, USA) were fixed on a vertical 1 m radius aluminum semi-circle frame with 0° elevation halfway along the arc. From -30° to 70°, we place 11 speakers with 10° separation and four at -80°, -60°, -45° and 90°. We used a 1/8” microphone (GRAS 46DP-1, GRAS Sound & Vibration, Denmark) fitted with a thin narrowing plastic tube and a 3D printed cuff as an acoustic coupler to measure sound in the ear of the bat. The opening of the thin end of the tube was inserted approximately at location of the tympanic membrane. The bat with the inserted microphone was placed a rotating, thin metal rod. The 360° rotation was controlled by a micro-controller (Raspberry Pi Pico RP2040, Raspberry Pi, United Kingdom) in steps of 5°. With the help of a two-plane laser we aligned the center of the speaker at elevation θ = 0° and θ = +90° with the turntable making sure all speakers had the same radial distance from the microphone.

Each speaker emitted a multi-harmonic stimulus (130 kHz to 15 kHz) (Supplemental Fig. A) with the help a custom driver (Supplemental Fig. B) and recorded the sound with the in-ear microphone with an UltraSoundGate 1216H from Avisoft (Germany, 375 kHz sampling rate, 16 bits). Last, we obtained a reference recording from all 15 speakers without the bat. We measured sound at a total of 1080 points (15 × 360°/5°) and calculated the minimum phase filter given the difference-spectrum of the stimulus of the in-ear recording and its reference resulting in the individual Head-Related Impulse Response (HRIR). For comparison with the photogrammetry method, we then filtered sinusoids at relevant frequencies with the measured HRIR to get the frequency specific HRTF.

## 3. Results

In total we created photogrammetry models of two deceased *M. daubentonii* (*Mdau 1* and *Mdau 2*), two deceased *M. nattereri* (*Mnat 1* and *Mnat 2*), one 3D printed *M. daubentonii*, and one living *M. daubentonii*. Of these we created μCT models of *Mdau 1* and *Mdau 2* and the 3D printed *M. daubentonii* and measured HRTF using an in-ear microphone and surrounding speakers from *Mdau 2, Mnat 1*, and *Mnat 2*.

### 3.1 Photogrammetry and μCT

We directly compared the aligned photogrammetry and μCT mesh models of the 3D print by computing the signed distance from the photogrammetry model to the μCT model (Fig. 2a). A per-vertex distance of 0 μm indicated the two models occupy the same geometric space, while a positive or negative per-vertex distance indicated the photogrammetry model is raised or lowered relative to the μCT model at that vertex. The comparison shows that more than 71.3% of vertices in the photogrammetry model have an absolute distance to the μCT scan of less than 50 μm (Fig. 2a). Fewer than 6.6% of all vertices have an associated distance larger than 150 μm mainly located inside the ear canal, where line-of-sight was compromised for the photogrammetry.

**Figure 2:**
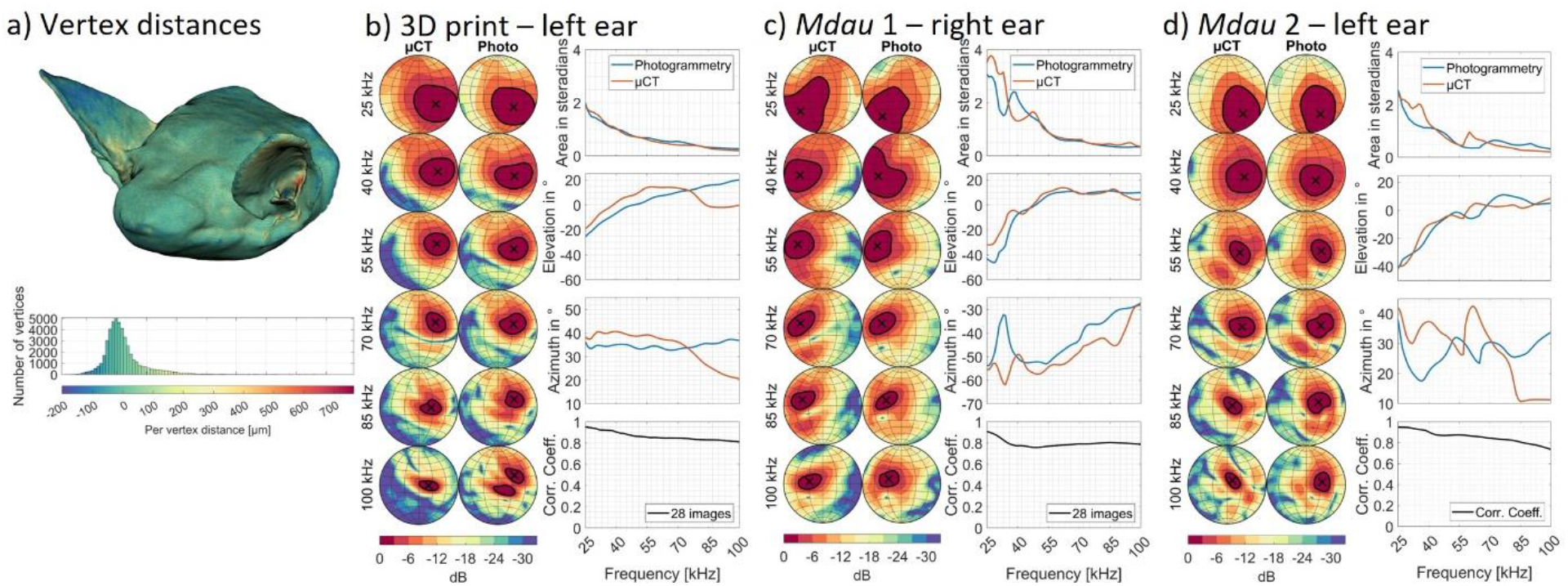
Comparing μCT and photogrammetry for model generation and HRTF simulations. a) Mesh model with color coded per-vertex distances from photogrammetry model to μCT model with the distribution of the vertex distances below. b-d) Frequency-specific, simulated HRTFs with region of highest sensitivity highlighted (−3dB from peak, black solid line, contour centroid, X). The area and centroid’s spatial orientation in azimuth and elevations angles of the region of highest sensitivity and correlation coefficients are plotted against frequency.

To compare different HRTFs, we calculated the correlation coefficient between the spatial information for the entire frequency range as in De Mey et al. (2008). The correlation coefficients for the HRTFs based on the μCT and photogrammetry models confirm a high degree agreement for all test cases (Fig. 2 b-d).

Despite the high per-vertex distances in the ear canal, the simulated HRTFs for the 3D printed bat show very good overall correspondence for all frequencies with deviations in elevation and azimuth at 75 kHz and above (Fig. 2b). Likewise, the two deceased *M. daubentonii* (Figs. 2 c-d) show good correspondence for all frequencies with larger discrepancies for the azimuthal location of the main lobe for *Mdau 2*.

### 3.2 Photogrammetry and measured HRTF

Comparing the photogrammetry-based simulated HRTFs with traditional HRTF measurements using an in-ear microphone again shows very good agreement for all four animals (Fig. 3). All correlation coefficients confirm a high degree of similarity between the simulated and measured HRTFs. The area, elevation and azimuth measurements of the main lobe all follow very similar patterns across frequencies with minor deviations except a large disagreement for the azimuthal location of *Mdau 2*).

**Figure 3:**
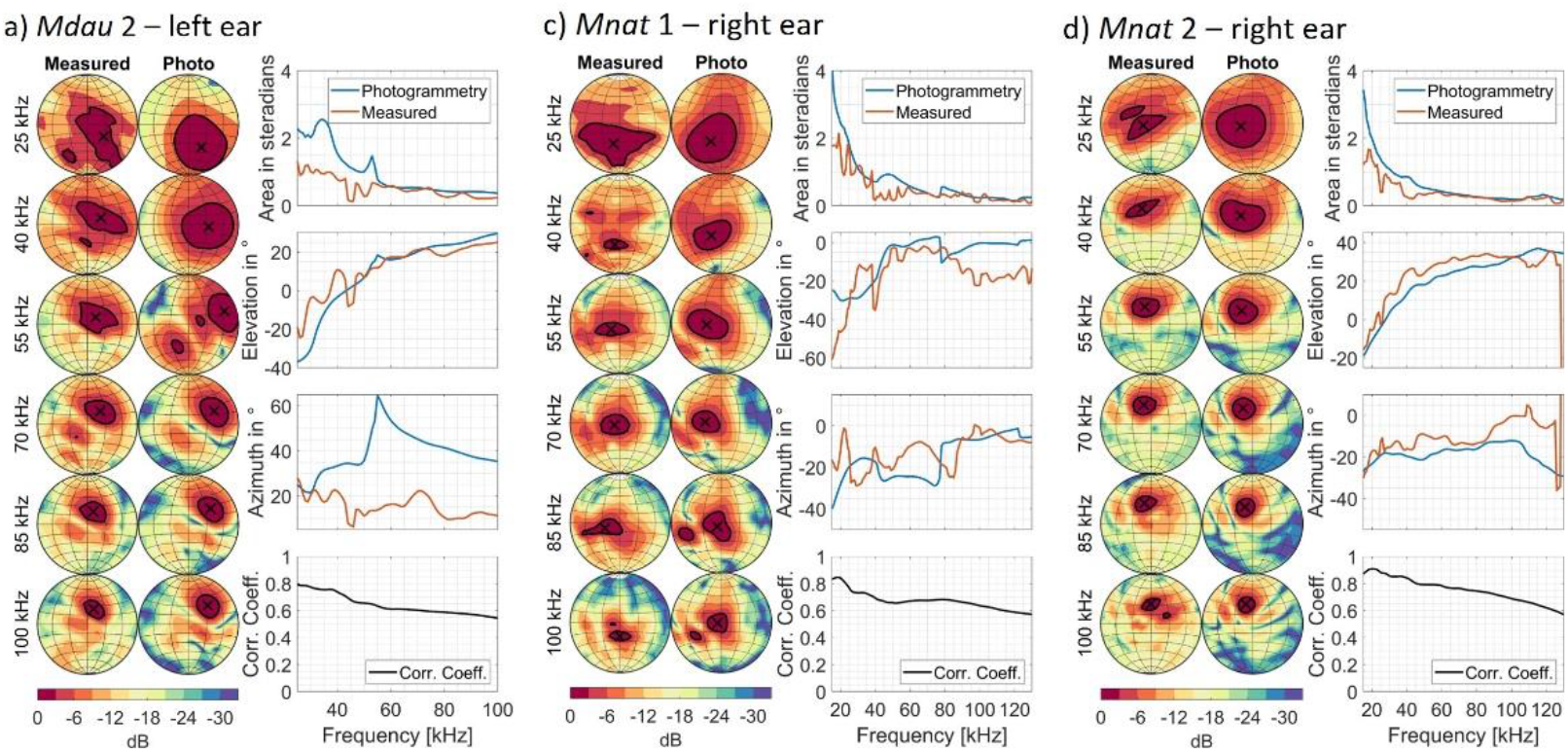
Measured HRTFs compared with simulated HRTFs of the 3D models generated with photogrammetry (a-c). Frequency-specific, simulated HRTFs with region of highest sensitivity highlighted (−3dB from peak, black solid line, contour centroid, X). The area and centroid’s spatial orientation in azimuth and elevations angles of the region of highest sensitivity and correlation coefficients are plotted against frequency.

### 3.3 Live animal photogrammetry

Finally, we photographed a live and awake *M. daubentonii* using the 28-camera setup. From the 28 captured images we generated a 3D model and simulated the HRTF (Fig. 4). When comparing a sample image (Fig. 4a) with the resulting 3D model (Figs 4b), the shape of the right pinna and tragus was successfully reconstructed. The boundary on the inside of pinna of the 3D model followed the boundary visible in the images, including the ridges visible on the inside of pinna right above the tip of tragus. The photogrammetry model was able to nicely replicate and separate the structural components deep into the pinna up until where the ear canal starts. Only a few intricate parts below tragus that have not been reconstructed successfully. The region of highest sensitivity has an exponentially decaying area as frequency increases and the elevation and azimuth vary smoothly with frequency. These features compare well with other simulated HRTFs presented in this study.

**Figure 4:**
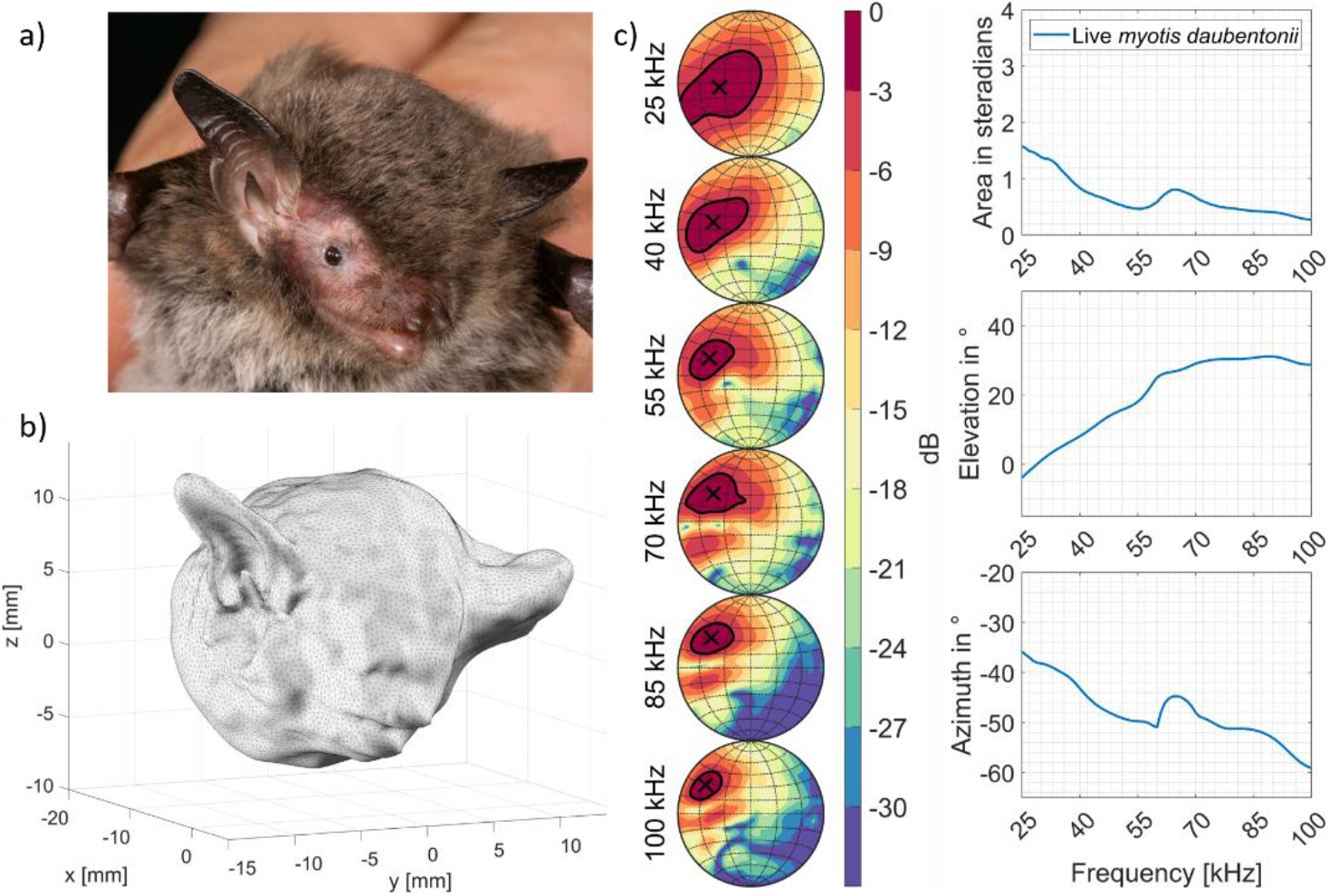
Photogrammetry and HRTF from an alive and awake *M. daubentonii*. a) One example image of the 28 used to generate 3D model through photogrammetry b) resulting 3D model with the simulated HRTF (c) including region of highest sensitivity (black solid line) and the area centroid (marked by X).

The physical size of the cameras including lenses limits both the number of available images and the angular separation between each image in the 28-camera setup. We investigated how these two factors affect the 3D models and subsequent acoustic modelling using the 3D print and *Mdau 1* as test cases. From the 511 images used in the previous analysis employing the 7-camera turntable rig (Fig. 1a), we selected 28 images that adhered to the spatial configuration of the 28-camera rig and used these to generate 3D models and simulated the HRTFs of the left ear of the 3D print and the right ear of *Mdau 1* (Fig. 5). The generated 3D models clearly loose detail especially of the opposing ear, but for both the 3D print and *Mdau 1*, the resulting HRTFs are strikingly similar to the full image set (lowest correlation coefficient of 0.87 for the 3D print and 0.66 for *Mdau 1*).

**Figure 5:**
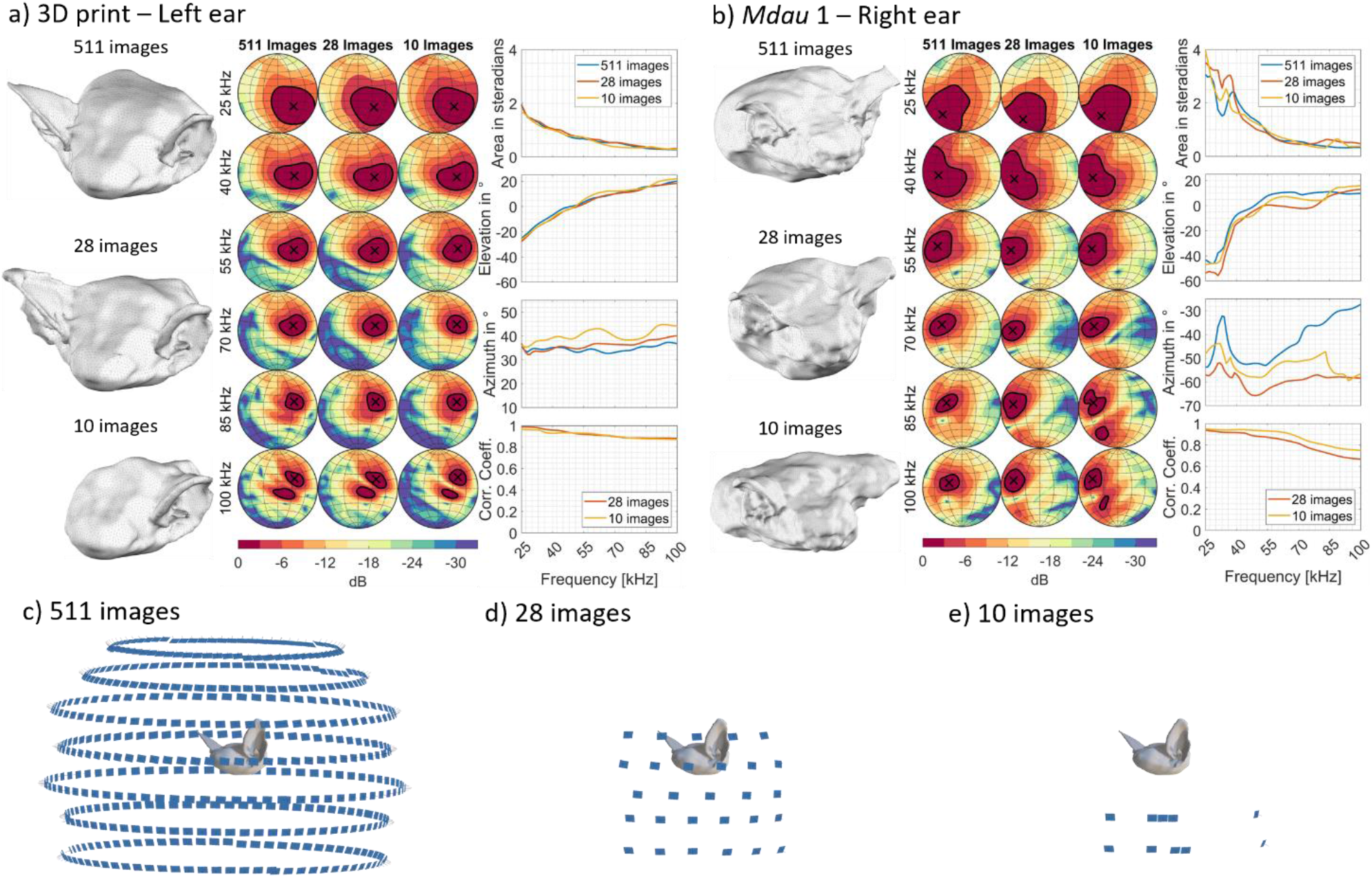
Investigating effects of the number of images and their spatial configuration on both the reconstructed 3D model and the simulated HRTF for two test cases: a) 3D print and b) *Mdau 1*. c) Spatial configuration of the 511 images created in the 7-camera turntable setup. d) Spatial configuration of images imitating a 28-camera rig for live animals and e) minimal number and unrestricted spatial configuration of images needed to reconstruct 3D model and successfully model the HRTF.

We also tested the minimum number of images necessary to simulate HRTFs if we ignore camera placement restrictions. We found that 10 images would be sufficient to generate a scaled 3D model and simulate the HRTF. The 10 images cover ca. 75° in azimuth and ca. 15° in elevation with some images being only 5° apart (see Fig. 5e). The similarity of the resulting HRTFs is in fact higher for the 10 image models of *Mdau 1* when comparing the correlation coefficients in Fig. 5.

## 4. Discussion

Our study clearly shows that photogrammetry can generate 3D models accurate enough to numerically model the HRTF of small animals at biologically relevant frequencies. When comparing our photogrammetry model directly with μCT models, the absolute difference is very small and errors larger than 50 μm only arise in locations where the camera view is limited. While this does create errors deep within the ear canal, both the models for the 3D print and of the deceased animals, show that these errors have little to no effect on the composition of the HRTF. Likewise, comparing photogrammetry with experimentally measured HRTFs show very little difference between these two approaches confirming the overall applicability of BEM and the accuracy of the 3D models. Photogrammetry is clearly a valid and cheap alternative to μCT scans for generating 3D surface models, even when violating good photogrammetry practice of having large angles (∼90°) between cameras that have line-of-sight of the same point on the subject. Furthermore, we show that it is possible to obtain 3D models suited for acoustic modelling using 28 images or fewer with our setup and pipeline, showing that it is entirely possible to obtain accurate models with a one-shot setup. In fact, we can accurately model the HRTF from as few as 10 images by only reconstructing one pinna, confirming previous results showing that for bats the head and body contributes little to the overall HRTF (De Mey, et al., 2008; Müller, 2004; Müller, et al., 2006). Thus, we clearly show that using photogrammetry, we can obtain accurate HRTF measurements even from alive and awake bats.

Our method of combining photogrammetry and acoustic modelling has several far-reaching implications: i) it allows for rapid, large-scale acquisition of images for HRTF simulation facilitating larger sample sizes than previously attainable. ii) It allows for none-invasive measurements, reducing or even abolishing the need for euthanasia and anesthesia. This expands the application of our method to animals where termination and/or the risk of anesthesia is not an option, e.g. studies of hearing development over time or measurements from highly endangered/protected species. iii) It will reliably provide realistic pinna orientation and shape for alive and awake subjects. Both freezing and fixation in ethanol affect the tissue properties and to a large degree also orientation and shape of the pinna. Using awake subjects ensures that the tissue is not compromised, and that the configuration is natural. However, it is noteworthy that the handheld placement of the awake subject may not reflect the in-flight/unrestrained configuration.

Photogrammetry is already widely used for morphological measurements (Irschick et al., 2022) and our results show that the 3D model accuracy is sufficient to allow for accurate numerical modeling of HRTFs from small animals at high frequencies as well. This also means that the method is easily applicable to larger subjects including humans assuming that the pinna morphology allows line of sight for all important features (pinna shape, orientation, ridges etc.). One caveat is reconstruction of hair; while the acoustic impedance of hair is ignored in our implementation of BEM as is done by DeMey et al. (2008), 3D reconstruction using photogrammetry creates large unrealist structures when hair is present. This means that animals such as foxes and cats with fur in their pinna will be difficult to model unless the fur is physically removed beforehand. The possibility of obtaining models from awake animals using photogrammetry makes this approach compared to other 3D scanning techniques attractive to many fields beyond acoustic modelling due to the here shown 3D model accuracy and affordability of the method itself.

While measured HRTFs represent the golden standard, modeled HRTFs inherently come with several advantages. Both the spatial resolution, and in particular the frequency range of the measured HRTFs is limited by available equipment (stimulus creating speaker and recording microphone). This means that it becomes exceedingly difficult to measure biologically relevant HRTFs from very high-frequency animals such as Percival’s trident bat, *Cleotis Percivali* (Fenton & Bell, 1981) with echolocation calls >150 kHz. The frequency response in the simulation is only limited by the size of the mesh elements in the 3D model, and for the frequency ranges and 3D models presented in this study, this limit was not reached. 3D models generated by μCT, or photogrammetry also enable structural modifications to pinna, tragus, or the entire head, to investigate what aspects of the morphology affects the HRTF without permanently destroying the original specimen. Moreover, modifications to the 3D model can far exceed the original morphology, e.g. extending the size of the pinna or tragus providing unprecedented insight into their functionality. The advantages of exploring form and function using such “digital surgery” extends far beyond acoustic modelling and we expect that the ease of generating 3D models with micro-meter precision using photogrammetry will prove a valuable tool for gaining scientific insight across multiple research fields.

## Acknowledgements

The study was funded by a Villum Fonden young investigator grant (00025380) to L.J. We are grateful for the guidance on photogrammetry of living animals given by Duncan Irschick. We give thanks to the machine workshop at SDU, especially Mikkel Stolberg Nielsen for building the camera rig. We also give thanks Danuta Wisniewska from our lab has been valuable in discussions throughout the project.

## Supplemental information

**Table S1:** All software used in the present study

| Software | Version | Description | Source |
| --- | --- | --- | --- |
| MATLAB | R2024a | Numeric computing environment | mathworks.com |
| TriggerBoxController | 1.6 | Controller software for the TriggerBox hardware | support.esperhq.com |
| CaptureGRID 4 | 4.27 | Photography workflow application for tethered shooting and advanced camera control | kuvacode.com |
| XnConvert | 1.98 | Batch image converter | xnconvert.com |
| Agisoft Metashape Professional | 2.10 | Photogrammetric processing of images for generation of 3D models | agisoft.com |
| Amira | 5.2.1 | CT image visualization and analysis | amira.com |
| 3D Builder | 20.0.4.0 | 3D design tool for editing and repairing 3D models. | Apps.microsoft.com |
| Meshlab | 2023.12d | Software for processing and editing 3D meshes. | Meshlab.net |
| Autodesk Meshmixer | 3.5.474 | Software for processing and editing 3D meshes. | Meshmixer.com |
| Blender | 3.6.2 | 3D creation suite that supports the entire 3D pipeline. | Blender.org |
| Mesh2hrtf | 1.1.1 | Toolbox for Numerical computation of head-related transfer functions. | Mesh2hrtf.org |

**Table S2:**
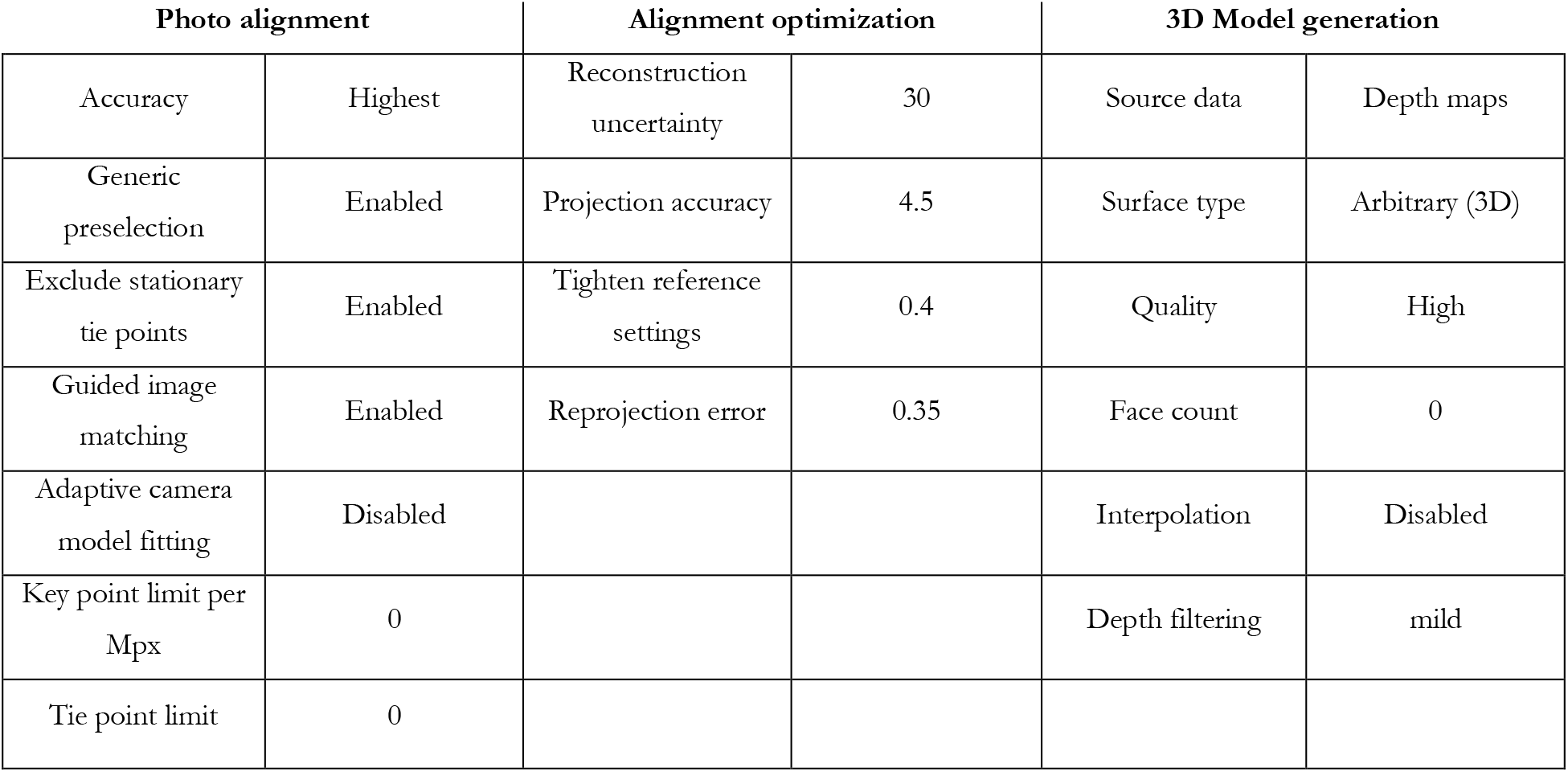
Parameters for generating photogrammetry models in Metashape

**Figure A:**
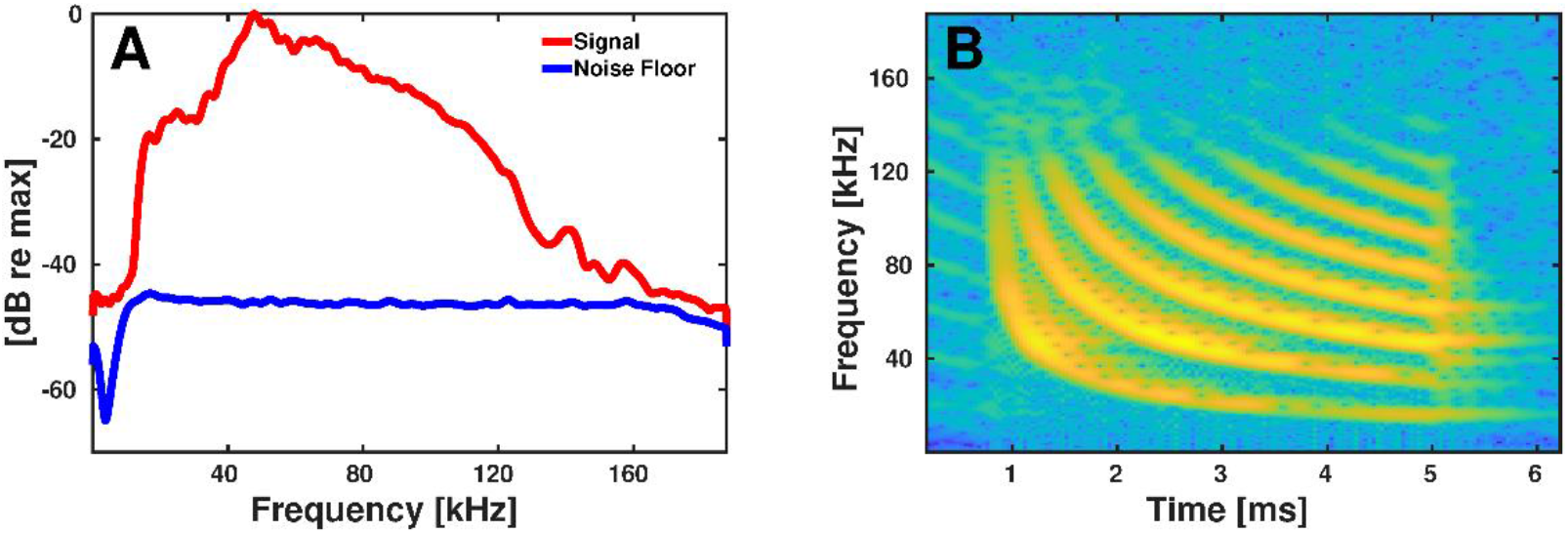
Multi-harmonic stimulus emitted for the measured HRTFs.

**Figure B:**
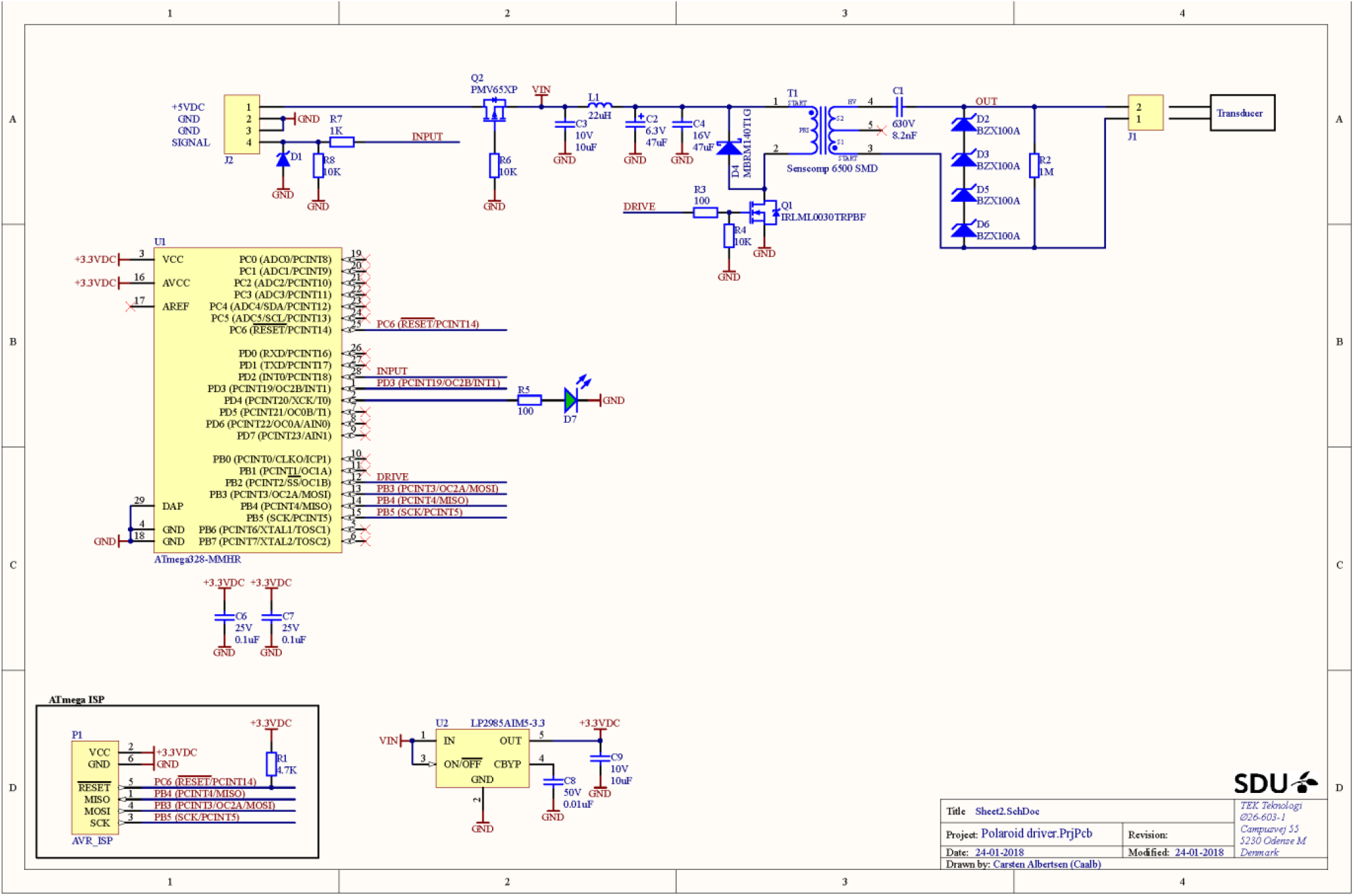
Schematic of the custom driver for the speakers used for the measured HRTFs.

